# Neural bases of proactive and predictive processing of meaningful sub-word units in speech comprehension

**DOI:** 10.1101/2024.04.29.591610

**Authors:** Suhail Matar, Alec Marantz

## Abstract

To comprehend speech, human brains identify meaningful units in the speech stream. But whereas the English ‘*She believed him.*’ has 3 words, the Arabic equivalent ‘*ṣaddaqathu.*’ is a single word with 3 meaningful sub-word units, called morphemes: a verb stem (‘*ṣaddaqa*’), a subject suffix (‘-*t*-’), and a direct object pronoun (‘-*hu*’). It remains unclear whether and how the brain processes morphemes, above and beyond other language units, during speech comprehension. Here, we propose and test hierarchically-nested encoding models of speech comprehension: a NAÏVE model with word-, syllable-, and sound-level information; a BOTTOM-UP model with additional morpheme boundary information; and PREDICTIVE models that process morphemes before these boundaries. We recorded magnetoencephalography (MEG) data as participants listened to Arabic sentences like ‘*ṣaddaqathu.*’. A temporal response function (TRF) analysis revealed that in temporal and left inferior frontal regions PREDICTIVE models outperform the BOTTOM-UP model, which outperforms the NAÏVE model. Moreover, verb stems were either length-AMBIGUOUS (e.g., ‘*ṣaddaqa*’ could initially be mistaken for the shorter stem ‘*ṣadda*’=‘*blocked*’) or length-UNAMBIGUOUS (e.g., ‘*qayyama*’=‘*evaluated*’ cannot be mistaken for a shorter stem), but shared a uniqueness point, at which stem identity is fully disambiguated. Evoked analyses revealed differences between conditions before the uniqueness point, suggesting that, rather than await disambiguation, the brain employs PROACTIVE PREDICTIVE strategies, processing the accumulated input as soon as any possible stem is identifiable, even if not unique. These findings highlight the role of morpheme processing in speech comprehension, and the importance of including morpheme-level information in neural and computational models of speech comprehension.

**Significance statement:** Many leading models of speech comprehension include information about words, syllables and sounds. But languages vary considerably in the amount of meaning packed into word units. This work proposes speech comprehension models with information about meaningful sub-word units, called morphemes (e.g., ‘*bake-*’ and ‘*-ing*’ in ‘*baking*’), and shows that they explain significantly more neural activity than models without morpheme information. We also show how the brain predictively processes morphemic information. These findings highlight the role of morphemes in speech comprehension and emphasize the contributions of morpheme-level information-theoretic metrics, like surprisal and entropy. Our models can be used to update current neural, cognitive, and computational models of speech comprehension, and constitute a step towards refining those models for naturalistic, connected speech.

## Introduction

Human speech unfolds over time, often as a continuous, connected stream of content at several hierarchical levels, such as sounds, syllables, and words. Behavioral and neural work has shown that in speech comprehension, the brain tracks speech sounds (called phonemes; Gwilliams et al., 2022), syllables (Saffran et al., 1996; Pefkou et al., 2017), and words (Cunillera et al., 2009). Moreover, leading neural (Hickok and Poeppel, 2007) and computational (McClelland and Elman, 1986; Norris and McQueen, 2008; Magnuson et al., 2020) models of speech comprehension account for words, syllables, and sounds.

To comprehend any sentence, the brain must retrieve its meaning. Of the three levels mentioned, only words carry meaning. But the notion of a ‘word’ varies considerably across languages (Dixon and Aikhenvald, 2003). For example, the English sentence ‘*She believed him.*’ has 3 meaningful, combinable words. But the Arabic equivalent ‘*ṣaddaqathu.*’ is a single, complex word that word-level processing cannot break down further. Thus, alone, word-level processing is limited: to comprehend Arabic, the brain would map entire unsegmentable sentences to meanings (‘*She-believed-him.*’) that are independently stored for all subject-object combinations (e.g., ‘*We-believed-her.*’, ‘*They-believed-me.*’, etc.). Considering how this scales for all other verbs and structures, Arabic word-level processing becomes computationally much costlier than English.

But words are not atomic units of meaning. ‘*Ṣaddaqathu.*’ has 3 meaningful sub-word units, called *morphemes*: ‘*ṣaddaqa-*’ (‘*believed*’) is a verb *stem*; ‘*-t-*’ (‘*she*’) is a subject-marking *suffix*; ‘*-hu*’ (‘*him*’) is a direct object pronoun (also called a *clitic*). Morpheme-level processing reduces complexity differences across languages (Creutz, 2006).

Past work supports a mental lexicon organized around morphemes (Marslen-Wilson et al., 1994). Additionally, in reading comprehension, the brain decomposes words into morphemes (Zweig and Pylkkänen, 2009; Solomyak and Marantz, 2010; Gwilliams and Marantz, 2015). Though fewer studies have tackled auditory morpheme processing, these typically find bilateral temporal and inferior frontal effects (Tyler et al., 2005; Bozic et al., 2010, 2015; Szlachta et al., 2012) within 300 ms of critical points (Leminen et al., 2011, 2013; Whiting et al., 2013; see Leminen et al., 2019, for a review).

Here, we ask *how* the brain processes spoken morphemes, and whether that is distinct from word-, syllable-, or phoneme-level processing. Specifically, is morpheme processing PREDICTIVE (ahead of morpheme boundaries) like other speech levels (Klimovich-Gray et al., 2019) or strictly BOTTOM-UP (post-boundary)?

The brain may predictively process spoken morphemes because they are often identifiable before unfolding fully. For example, the spoken English word ‘*bargain-ed*’ (phonetically: /bɑrgənd/) has a stem that is *uniquely* identifiable already during the /g/ sound: ‘*bargain*’ is the only English morpheme with the onset sequence /bɑrg/. Past work shows *stem uniqueness* points are behaviorally relevant to spoken word recognition (Balling and Baayen, 2008). But there could be another, earlier predictive point: /r/ is the *first stem identification* point, at which a (wrong) stem (‘*bar*’) first becomes identifiable.

We built several hierarchically-nested encoding models of speech comprehension. We also propose 3 predictive strategies for spoken morpheme processing: (i) a PATIENT strategy, where the brain awaits *stem uniqueness* (/bɑrg/) before fully identifying morphemes; (ii) an EAGER strategy, where the brain deterministically processes the first identifiable stem (at /bɑr/), even if not unique, then corrects its error if necessary as more input accumulates; (iii) a PROBABILISTIC strategy, where the brain hedges its bets: at the *first stem* point it uses distributional information to assign different weights to a short (‘*bar*’) or long (‘*bark*’, ‘*bargain*’, etc.) stem, then shifts those weights as more input accumulates. The latter two are PROACTIVE strategies, processing the accumulating input before full disambiguation.

To test these models, we recorded cortical activity using magnetoencephalography (MEG) as native Arabic speakers listened to multi-morphemic sentences such as ‘*ṣaddaqa-t-hu.*’ (‘*She believed him.*’). Our design capitalized on Arabic’s grammatical properties, and allowed for two complementary analyses: Temporal Response Function (TRF) analyses to compare pairs of models as to their goodness-of-fit to the neural data; and evoked analyses using an embedded categorical design.

## Materials and Methods

### Participants

We recruited 28 native Arabic speakers to participate in the experiment at New York University (NYU). The NYU Institutional Review Board approved the experiment, which we performed in accordance with the relevant regulations. Participants had typical hearing, and normal or corrected-to-normal vision. They all provided written consent and received compensation for their time. We excluded one participant’s data due to a poor behavioral score (at chance in one of the two types of tasks; see *Experimental Task*). Of the remaining 27 participants (mean age=29.9, SD=3.7), 16 were female.

### Experimental Design

Participants listened to single-word multi-morphemic sentences (e.g., ‘*ṣaddaqa*-*t*-*hu*’, meaning ‘*She believed him.*’) in Modern Standard Arabic (Fig. 1a). Each item was a full sentence consisting of a verb stem morpheme (‘*ṣaddaqa*’=‘*believed*’), a subject suffix (‘*-t-*’, indicating ‘*she*’ as the subject) and a direct object pronoun (‘*-hu*’, indicating ‘*him*’ as the object).

**Figure 1.**
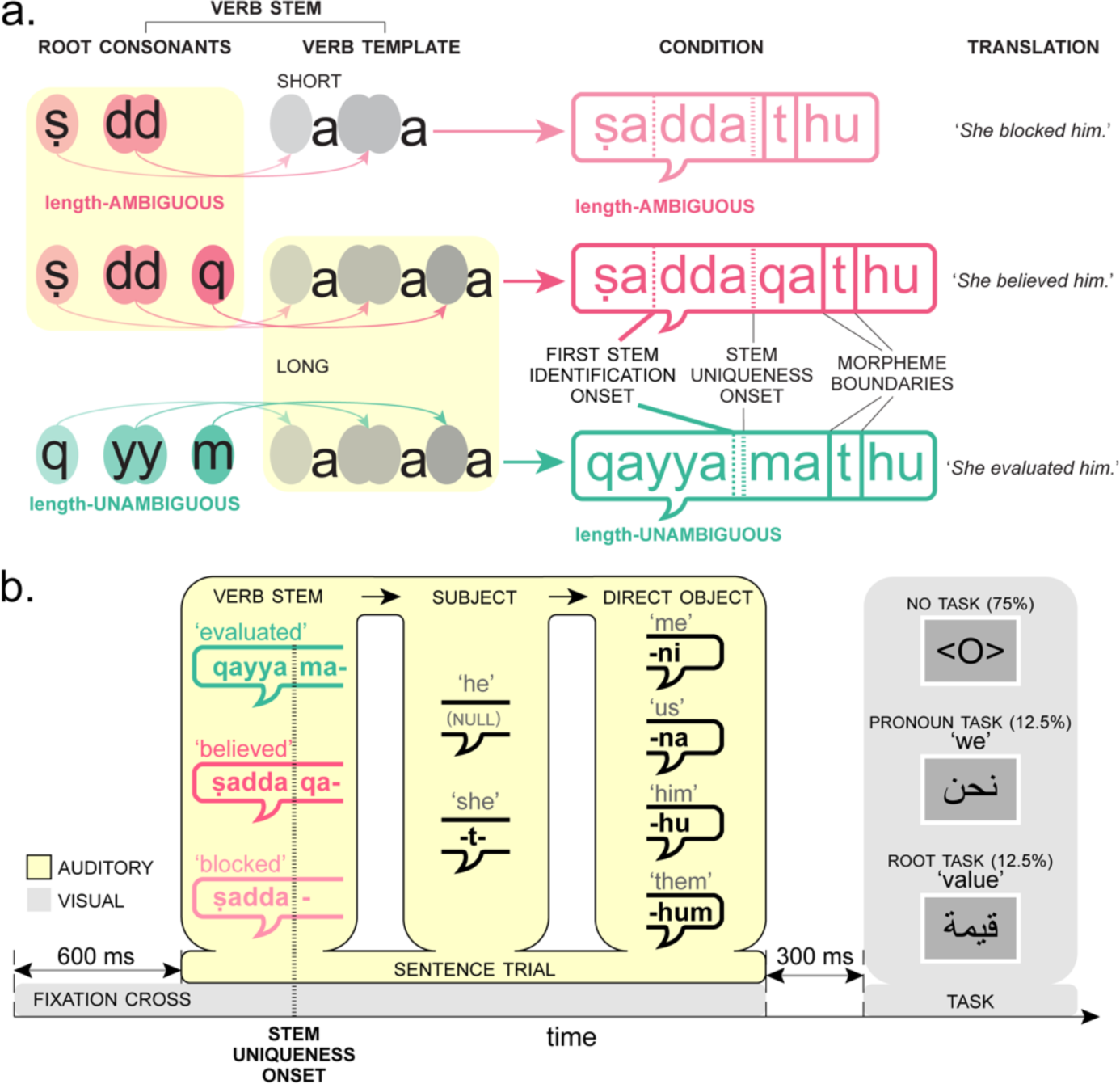
Study design and trial structure. (a) Stimuli were multi-morphemic words whose first part is a verb stem. Stems shared one of two Arabic vocalic verb templates: short (with 2 consonant placeholders; gray ovals) or long (3 consonant placeholders); the short template is identical to the onset of the long template. Stems belonged to two conditions. Temporarily length-_AMBIGUOUS_ stems (pink) were pairs of long and short stems, where the short stem (‘*ṣadda*’) was identical to the long stem’s onset (‘*ṣaddaqa*’). Length-_UNAMBIGUOUS_ stems (turquoise) were such that their onsets could not be confused with a shorter stem in the language (‘*qayyama*’). All stems shared a *stem uniqueness onset* (dashed line)*—*the point after which the stem is uniquely recognized among all stems in the language. But the *first stem identification onset* (dotted line)—at which a stem is first identifiable in the input— differed across conditions. In length-_UNAMBIGUOUS_ stems, this point was identical to *stem uniqueness onset*. In length-_AMBIGUOUS_ stems, the *first stem identification onset* was earlier. (b) Trial structure. Each trial began with a visual fixation cross on screen for 600 ms, at which point the auditory trial began. Each stimulus consisted of a multi-morphemic word that formed a full sentence with a verb stem morpheme, a subject morpheme (1 of 2), and a direct object morpheme (1 of 4). 300 ms after audio offset a visual task screen appeared.

We used 90 verb stems, pairing each with 8 combinations of 2 subject suffixes and 4 direct object morphemes (Fig. 1b), resulting in 720 items. Subject suffixes were either ‘*-t-*’ (‘*she*’) or a null morpheme (‘*he*’). Direct object pronouns were ‘*-ni*’, ‘*-na*’, ‘*-hu*’, and ‘*-hum*’ (corresponding to ‘*me*’, ‘*us*’, ‘*him*’ and masculine ‘*them*’, respectively; Arabic has 12 such pronouns). Each verb stem has its own probability distribution over which direct object pronouns follow. If the brain predictively processes morphemic information during comprehension, then information theory metrics derived from these distributions (see *Cognitive models*) should explain cortical activity.

Verb stems were either long or short (Fig. 1a). Long stems consisted of 3 root consonants {***C_1_***, ***C_2_***, ***C_3_***} substituting placeholders in a long verb template ***C_1_****a****C_2_C_2_****a****C_3_****a*. For example, ‘***ṣ****a**dd**a**q**a*’ (‘*believed*’) and ‘***q****a**yy**a**m**a*’ (‘*evaluated*’) have this template, with respective root consonants {**ṣ**, **d**, **q**} and {**q**, **y**, **m**}. Short stems (e.g., ‘***ṣ****a**dd**a*’=‘*blocked*’) consisted of 2 root consonants {***C_1_***, ***C_2_***} in a short verb template, ***C_1_****a****C_2_C_2_****a*. The short template is identical to the onset of the long template. All long stems and all short stems shared their respective templates.

Each short stem (e.g., ‘***ṣ****a**dd**a*’) had a corresponding long stem (‘***ṣ****a**dd**a**q**a*’), with which it shared its two consonants ({**ṣ**, **d**}; Fig. 1a), making temporarily length-AMBIGUOUS pairs (similar to the *bar*–*bargain* example; see *Introduction*). However, half the long stems were length-UNAMBIGUOUS (e.g., ‘***q****a**yy**a**m**a*’=‘*evaluated*’), since its onset does not form an Arabic stem (e.g., ‘***q****a**yy**a*’ is not a stem). Importantly, all verb stems across conditions had the same *stem uniqueness onset* (Fig. 1), after which the stems may be uniquely identified from all other stems in the language; this point was at the offset of the sequence *C_1_aC_2_C_2_a* (Fig. 1a). For example, before the onset of /m/ in ‘*qayya**m**a*’, the competing stem ‘*qayya**d**a*’ (‘*tied*’) is a viable candidate that can only be discarded after the onset of /m/.

But here we propose another potential decision point. We define the *first stem identification* point as the point at which a stem is first identifiable in the input (Fig. 1a). For length-AMBIGUOUS items, this corresponds to ***C_2_*** onset—the first point at which a viable root consonant set (with 2 consonants; e.g., {**ṣ**, **d**}) and a stem (‘***ṣ****a**dd**a*’) are first identifiable, even if erroneously. But for UNAMBIGUOUS items, the *first stem identification* onset is identical to *stem uniqueness* onset—only as of that point is a viable root available (now, with 3 consonants; e.g., {**q**, **y**, **m**}).

In sum, we used three types of stems: 30 short length-AMBIGUOUS stems, 30 long length-AMBIGUOUS stems, and 30 length-UNAMBIGUOUS stems (see Supplementary Table 1 for full list of stems).

We use this design in complementary encoding (TRF) and evoked analyses to test 3 different PREDICTIVE parsing strategies. A PATIENT parser is fully risk-averse; it treats all trials similarly, waiting until *stem uniqueness onset* to fully disambiguate and identify the stem morpheme; this parser is oblivious to the *first stem identification* point (Fig. 2). An EAGER parser responds deterministically to the *first stem identification* point, even at the cost of later error correction. A PROBABILISTIC parser responds probabilistically to the *first stem identification* point, assigning weights to different stem-length possibilities based on language statistics. The EAGER and PROBABILISTIC parsers differentiate between the two conditions: only in length-AMBIGUOUS stems is the *first stem identification* distinct from *stem uniqueness* (Fig. 1a & 2).

**Figure 2.**
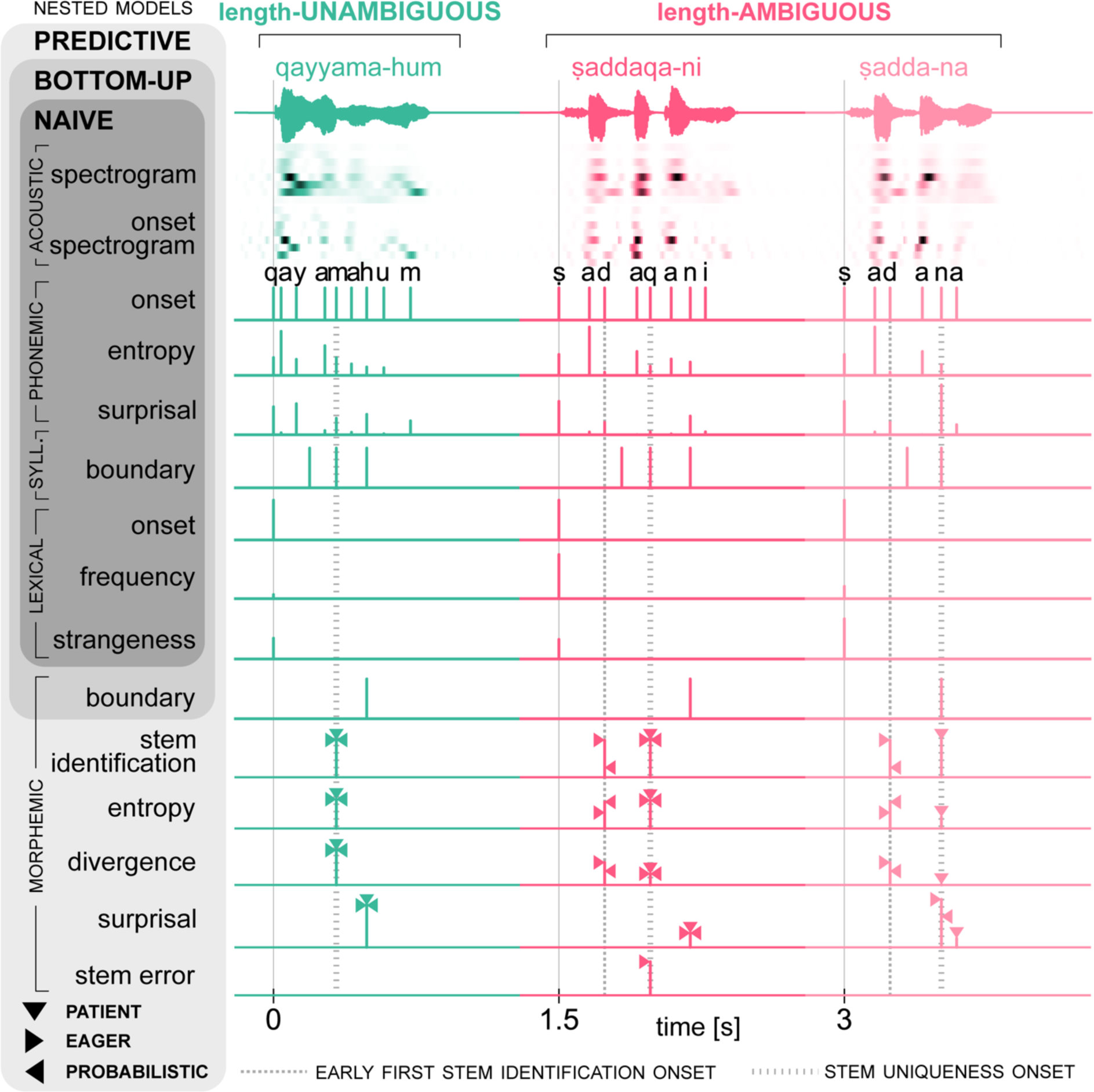
Encoding models of morpheme processing in speech comprehension. We propose 3 types of hierarchically-nested encoding models. A _NAÏVE_ model had acoustic, phonemic, syllabic, and lexical features, but no morphemic information. It was nested within a _BOTTOM_-_UP_ model that had an additional morpheme boundary feature. This model was nested inside _PREDICTIVE_ models, which had additional morpheme-based information-theory metrics. We built 3 _PREDICTIVE_ models which differed in the predictive strategy used: _PATIENT_ (▾), _EAGER_ (▶), or _PROBABILISTIC_ (◀). Each model contained a bundle (gray rectangle) of features. Apart from acoustic features, all features were impulse trains, whose values were 0s except for critical time-points (see Table 1 for definitions and timings). For _PREDICTIVE_ features, the value for each strategy is indicated by the triangular apex touching the impulse. The figure shows feature values for the 3 example stems in Fig. 1. Acoustic features are spectrograms calculated in 8 logarithmic frequency bands (100 Hz–5 kHz; see Materials and Methods).

**Table 1.** Impulse values and timings of each model feature.

| Feature: definition | Impulse value | Impulse timing |
| --- | --- | --- |
| **Phoneme onset** | 1 | Phoneme onset |
| **Phoneme entropy:** uncertainty regarding the next sound, given the current sound sequence $S$ | $-\sum_{x \in X} P(x\vert S) \cdot \log_2 P(x\vert S)$ | |
| **Phoneme surprisal:** how unlikely the current phoneme $x$ is, given the preceding sound sequence $S$ | $-\log_2 P(x\vert S)$ | |
| **Syllabic boundary** | 1 | Syllabic boundary |
| **Word onset** | 1 | Word onset |
| **Word frequency**<br>(van Heuven et al., 2014) | $\log_{10}(\text{frequency per billion words})$ | |
| **Word strangeness:** how lexically strange the combination of stem and direct object pronoun is (see *Experimental Design*) | From behavioral norming study |  |
| **Morpheme boundary** | 1 | Morpheme boundary |
| **Stem identification:**<br>PATIENT, EAGER: unit impulse upon stem identification.<br>PROBABILISTIC: probability of the current sequence $V$ being the stem (and not the onset of a longer stem). | PATIENT and EAGER: 1<br>PROBABILISTIC: $P(\text{stem} = V\vert V)$ | PATIENT: at *stem uniqueness onset* only<br>EAGER, PROBABILISTIC: at *stem uniqueness onset only* (length-UNAMBIGUOUS trials); at *first stem identification onset* and *stem uniqueness onset* (length-AMBIGUOUS trials) |
| **Morpheme entropy:** uncertainty over direct object pronouns, given the identified verb stem $V$ | $-\sum_{m \in M} P(m\vert V) \cdot \log_2 P(m\vert V)$ | |
| **Morpheme divergence:** compared to the probability distribution over direct object pronouns across all ($A$) verb stems in the language, how divergent is the probability distribution following the identified stem $V$ | $-\sum_{m \in M} P(m\vert V) \cdot \log_2 \left[ \frac{P(m\vert A)}{P(m\vert V)} \right]$ | |
| **Morpheme surprisal:** how unlikely the current morpheme $m$ is, given the identified stem $V$ | $-\log_2 P(m\vert V)$ | Direct object onset |
| **Stem error detection:** (EAGER only) indicator for error in the deterministic commitment to the wrong stem | 1 | *stem uniqueness onset* |

We generated a sound file for each item using Google’s Arabic Text-to-Speech engine (24kHz sampling rate), which uses a female-sounding voice. When debriefed about the experiment, participants reported not guessing that the speech is machine-generated. We manually manipulated the sound files using Praat (Boersma and Weenink, 2021) to make the acoustic onset identical within each set. For each length-UNAMBIGUOUS stem, we chose one of 8 items (counterbalanced across sets) and used its ***C_1_****a****C_2_C_2_****a****C_3_*** onset to replace the onset of the remaining 7 items. For length-AMBIGUOUS stems, we followed a similar logic for each set of 16 items (8 long and 8 short with the same onset). If the chosen item had a long stem, we used its ***C_1_****a****C_2_C_2_****a****C_3_*** onset to replace the onset of the other 7 long-stem items, and its ***C_1_****a****C_2_C_2_*** onset to replace the onset of all 8 short-stem items; if the item chosen had a short stem, we used its ***C_1_****a****C_2_C_2_*** onset to replace the onset of all other 15 items, then randomly picked a long item, extracted its now-altered ***C_1_****a****C_2_C_2_****a****C_3_*** sequence to replace the onset of the remaining 7 long-stem items. All this guaranteed that participants could not use the acoustic onset alone to guess whether the verb stem is long or short (in length-AMBIGUOUS trials), whether the subject suffix is feminine or masculine, or which direct object follows. Thus, any early effects we find can only be attributable to participants’ implicit knowledge about the statistics in the language. The resulting stimuli varied in length between 652–1092 ms. We manually identified and annotated all onset, splicing, replacement, and boundary points (between phonemes, syllables, or morphemes) as the nearest zero-crossing in the selected location on the oscillograms.

Some items were implausible or infrequent combinations of a verb stem and an object (e.g., ‘*sabbaba-ni*’, meaning ‘(He) caused me.’). We included these items in order to have a wide range of values of transition probabilities between stems and direct object pronouns. But to account for these items’ strangeness, we ran an online behavioral norming study via Gorilla. We divided all items into eight lists, such that each verb stem appeared only once per list. Each of 200 Arabic speakers (via Prolific) rated the items from one list on strangeness (1–5 Likert scale), resulting in 25 ratings per item. We chose to ask about strangeness, rather than well-formedness, because the latter is harder to capture in pithy terms in Arabic, and because more semantically anomalous items typically lead to stronger brain responses (Friederici et al., 1993). For each item, we used its average norming rating to build the word strangeness feature (see *Cognitive Models*, Fig. 2, and Table 1).

### Experimental Task

After 75% of trials, there was no task; the symbol ‘<O>’ appeared on screen and participants pressed one of two buttons to continue. After the remaining 25% of trials, visual task items appeared on the screen, consisting of single words (Fig. 1b). There were two task types, represented equally.

In the first type, one of four pronouns (equivalent to ‘*I*’, ‘*we*’, ‘*he*’, and ‘*they*’) appeared on screen and participants judged if they corresponded to the direct object pronoun they heard in the trial. For example, if they heard ‘*him*’, and saw the task item ‘*he*’, they had to press the Correct button.

In the second task type, nouns or adjectives appeared on the screen and participants judged if they shared the same Arabic root (meaning, the same consonant set in Fig. 1a) with the verb stem. For example, if the verb stem was ‘***ṣ****a**ww**a**r**a*’ (‘*photographed*’, root={**ṣ**, **w**, **r**}) and the task item ‘***ṣ****u**wr**a***’** (‘*photograph*’, same root), they had to press the Correct button.

For each set of 8 items that shared the same stem, one item had the first task type and another had the second task type. Across sets, we counterbalanced which direct object pronouns were associated with each task, and with correct or incorrect responses. We chose these tasks for their ease, and to ensure participants parsed both stems and pronouns.

### Information theory metrics

Typically, predictive processes in language are operationalized using information theory metrics. Here, we use 3 such metrics: (i) Entropy measures the uncertainty about upcoming information (e.g., a direct object morpheme) given the current context (e.g., the identified stem) (Wurm et al., 2006); higher entropy means the context is less predictive of upcoming information; (ii) Divergence (also known as the Kullback-Leibler divergence; Aurnhammer and Frank, 2019) measures how deviant a probability distribution is from a typical distribution. Here, we first calculated the typical probability distribution over direct object pronouns across all verb stems in Arabic. Then, for each verb stem used, we calculated its own probability distribution over direct object pronouns, and measured how divergent this distribution is from the typical one; higher divergence means the identified verb stem behaves less like the typical stem in the language; (iii) Surprisal quantifies how unlikely the present information is, given the preceding context (Balling and Baayen, 2012); higher surprisal means the current information is more unlikely.

### Encoding models of morpheme processing

We built a series of hierarchically-nested encoding models of speech comprehension. Each model comprises a set of features (Fig. 2). A feature is a time series that captures one facet of the input. Apart from the acoustic features, all features were impulse trains. For example, a word onset feature signals when a word begins: its value is 1 at timepoints corresponding to word onsets, but is 0 otherwise. Table 1 defines all impulse features used, and explains their impulse values and timings. We calculated most feature values and metrics based on the Arabic Gigaword corpus (v5.0). To get morpheme information, we parsed the corpus using an Arabic morphological analysis and disambiguation algorithm (MADAMIRA—Pasha et al., 2014), then converted the output into a lexicon to extract frequency information and probability distributions.

The base model we built, called the NAÏVE model, is blind to morphemic information, but includes input features at the level of acoustics, sounds, syllables, and words (Fig. 2). Acoustic features include a gammatone-filtered spectrogram of the auditory input (8 logarithmic frequency bands in 100 Hz–5 kHz; applying gammatone filters approximates cochlear and early neural auditory processes; Heeris, 2024), and onset spectrogram (the half-wave rectified temporal derivative of the gammatone spectrogram; Brodbeck et al., 2020). Sound (phonetic) features included phoneme onset, phoneme entropy, and phoneme surprisal; phoneme entropy and surprisal are defined as functions of the probability distribution of a phoneme given the phonemic context (Table 1); they account for predictive processes at the sound level. At the syllable level, we included syllabic boundary as a feature, and at the word level we included word onset, word frequency, and word strangeness (see *Experimental Design*).

We nested this model within a BOTTOM-UP model, which includes all NAÏVE features, but with one extra feature at the morpheme level: morpheme boundary (Fig. 2, Table 1). The BOTTOM-UP model can only process morphemes after encountering a morpheme boundary.

Next, we built several morphologically PREDICTIVE models, each with its own parsing strategy: PATIENT, EAGER, or PROBABILISTIC. Each PREDICTIVE model has all of the nested BOTTOM-UP features, but with additional morphemic features (Fig. 2, Table 1): stem identification, morpheme entropy, morpheme divergence, and morpheme surprisal. The EAGER strategy additionally has a stem error detection feature. The values and timings of these morpheme features differed across the PREDICTIVE models, as outlined in Table 1.

Correlations between impulse trains (excluding unit impulse trains, such as phoneme onset or syllable boundaries) appear in Supplementary Fig. S1. To avoid inflated correlation values (impulse trains are 0 in most time-points), and since predictors can differ in terms of timings and/or values, we calculated correlations in two, complementary ways. Fig. S1a shows correlations between average impulse values: we averaged each feature’s values within each trial and correlated all pairs of the resulting vectors. Fig. S1b combines information about impulse values and timings: for each pair of features, we correlated all the pairs of points for which at least one feature is non-zero; thus, stronger positive correlations indicate impulses that are similarly timed and have positively correlated values, whereas stronger negative correlations indicate features that either have different timings, or similar timings but negatively correlated values. Together, the two methods give a more holistic image of the similarities and differences between features.

### Stimuli presentation

We divided the stimuli into 8 blocks following a Latin square design. Each stem was represented only once per block, and each block had the same number of task items. For each participant, we randomized block order and trial order within blocks. To alleviate participants’ experience, each block was divided into two in the actual experiment, resulting in 16 short blocks.

The experiment began with instruction screens in Arabic, followed by a short practice. This ensured all participants received the same explanation and content. We then verbally ensured that participants understood the task before proceeding with the experimental trials.

Each trial began with a fixation cross appearing in the center of the screen for the duration of the trial (Fig. 1b). After 600 ms, a trigger signal started playing the auditory sentence. After audio ended, the fixation cross disappeared, leaving a 300 ms blank screen, after which the task screen appeared until participants pressed either button. To avoid entrainment to presentation frequency, a blank screen appeared after button press for a random duration between 300–500 ms, sampled uniformly. Task items and fixation crosses appeared as white text against a gray background.

To present stimuli, we used PsychoPy (version 2021.2.3). The experiment lasted ∼45 minutes. A projector relayed the image to a screen inside the Magnetically Shielded Room (MSR), which houses the MEG sensors.

### MEG data acquisition

Before the experiment, we used a hand-held FastSCAN laser scanner (Polhemus, VT, USA) to digitize participants’ head-shape, plus five marker points on the head (center, left, and right of forehead, and one anterior of each auditory canal) and three anatomical points (nasion and bilateral tragi). Inside the MSR, we placed marker coils on the digitized marker points to align the head with the MEG sensors and the scanned head-shape. Marker measurements obtained right before and right after the experiment measured and accounted for overall head movement. During the experiment, we acquired MEG data using a 157-channel axial gradiometer system (Kanazawa Institute of Technology, Kanazawa Japan), with a 1 kHz sampling rate and an online 0.1–200 Hz band-pass filter.

### Structural MRI acquisition

We collected structural Magnetic Resonance volumes from 22 participants using NYU’s 3T MAGNETOM Prisma (Siemens, Germany), and a 64-channel head coil (TR=2.4 s; 256 slices; 0.8 mm×0.8 mm×0.8 mm). Volumes were pre-processed using Freesurfer (Fischl, 2012) and manually corrected for segmentation errors.

### MEG data preprocessing

We noise-reduced data using the Continuously Adjusted Least Squares Method (CALM—Adachi et al., 2001; MEG160 software v2.004A — Yokogawa Electric Corporation and Eagle Technology Corporation, Tokyo, Japan), which eliminates noise recorded in reference MEG channels located in the MSR but away from the head. We processed the data using MNE-Python (v0.23; Gramfort et al., 2014). We first applied a 1–40 Hz band-pass filter to the data. The 1 Hz filter is crucial for cleaning our data, due to excessive magnetic noise from the surrounding urban environment. We then identified bad channels (saturated/excessively noisy; min=1, max=9, median=2.5). We corrected trigger timings by identifying speech onset for each trial, then split the data into 1.5 s epochs, including a 200 ms baseline period. We applied independent component analysis (ICA) to the epoched data. Because of ambient magnetic noise, we calculated a clean ICA solution by iteratively identifying ICA components that represent noise bursts, then removing any noisy epochs giving rise to these components just for the sake of recalculating the ICA solution in the next iteration. Total percentage of epochs excluded for ICA purposes was kept below 10%. In the last iteration, we identified and removed noise components related to known biological (heartbeats, eye blinks, eye movements) or system-specific sources, then applied the ICA solution to all epochs, including excluded epochs. This method guaranteed that the ICA solution is not governed by few noise bursts explaining large portions of the data. Next, we interpolated the data in bad channels based on the remaining channels, applied a baseline correction, then automatically excluded any epochs with absolute values exceeding 3000 fT. On average, we discarded 1.8% of trials (SD=1.1%). Finally, we calculated a noise covariance matrix using the baseline periods of all trials.

For co-registration, we aligned the head-shape and anatomical marker points to the MR-reconstructed head surface and created a Boundary Element Model of the reconstructed cortical surface, from which we derived a source-space mesh of 2,562 sources per hemisphere. For the 5 participants without MR volumes, we used the Freesurfer average brain instead (Fischl, 2012). We calculated a forward solution matrix from the cortical surface to the MEG sensors, then used it with the noise covariance matrix to calculate the inverse solution using minimum-norm estimates. We constrained dipoles to the direction orthogonal to the cortical surface, yielding a signed estimate of activation in the form of noise-normalized dynamic Statistical Parameter Maps (dSPM; Dale et al., 2000)

### Regions of Interest (ROIs)

We constrained our analyses to 4 ROIs on the cortical surface, chosen from Freesurfer’s Desikan-Killiany atlas-based parcellation: the bilateral inferior frontal cortices (combining the labels *pars opercularis*, *pars triangularis*, and *pars orbitalis*) and the bilateral superior and middle temporal cortices (combining the labels *superior temporal*, *middle temporal*, *transverse temporal*, and *banks sts*). We chose these ROIs because of their demonstrated involvement in auditory and speech processing, including sensitivity to morpheme properties and information theoretic metrics of speech (Tyler et al., 2005; Bozic et al., 2010, 2015; Gagnepain et al., 2012; Szlachta et al., 2012; Ettinger et al., 2014; Gwilliams and Marantz, 2015; Willems et al., 2016; Brodbeck et al., 2018; Bhaya-Grossman and Chang, 2022).

### Temporal Response Function (TRF) calculation and statistical analysis

We performed encoding analyses to calculate TRFs using the Eelbrain wrapper for MNE-python (Brodbeck et al., 2023). We first calculated the inverse solution per single trial (SNR=2), morphed all solutions from participants’ native MRI space to the shared Freesurfer average surface, then concatenated each participant’s epochs. We decimated the result to 100 Hz for computational efficiency. (Since we had band-pass filtered the data at 1–40 Hz, this did not introduce aliasing concerns.) For each cortical source of each subject, the result was a 100 Hz time-course of cortical responses to all trials. This time-course was the dependent variable in all TRF analyses. We estimated TRFs using a mass-univariate approach, calculating TRFs independently for each model, for each subject, at each cortical source.

We estimated TRFs for all features jointly. Each feature’s TRF was built over time-lags of 0–500 ms; this means that we approximate brain activity at each timepoint by integrating information present in the preceding 500 ms of input. To estimate the TRFs, we used the boosting algorithm (David et al., 2007), which enforces a sparse solution that avoids overfitting the data. In each iteration, the algorithm adds a small ‘boost’ to one feature’s TRF, at the precise time-lag that results in the largest reduction in *l1* error between the neural time-course and the TRF-predicted activity. The boosts had a 50 ms Hamming kernel; note that this means that boosting a certain time-lag also changes five other surrounding time-lags. Thus, to keep the TRF solutions causal (the output predicted only by the past, not the future) the first boost-able point was at time-lag 20 ms. We also used selective stopping, freezing a feature’s TRF if boosting it further no longer caused a reduction in error, and continuing with the remaining features until they were all frozen.

In order to further avoid overfitting, we split the data into 5 partitions: we trained and validated the TRF solution using 4 partitions, and tested it on the remaining partition. We repeated the process 5 times; each partition was used for testing in one repetition. In each repetition we performed 4 runs, where we alternated which of the 4 non-test partitions was the validation partition, resulting in 20 TRF solutions (5 folds × 4 runs). The algorithm iteratively built each TRF solution using the training and validation partitions. Later, it averaged the TRF solutions across the 4 runs. We convolved each partition’s encoding model with the average TRF solution across the 4 corresponding runs, resulting in the model’s *prediction* of the real neural data in the held-out test partition. Finally, we concatenated all predicted partitions and compared the result against the real data, calculating the residual gap between the two.

Repeating this process for every cortical source resulted in a cortical map of explanatory power for each subject, for each model. In our analyses, we perform group-level comparisons between explanatory power maps of pairs of models, since our models are hierarchically-nested. The null hypothesis in these comparisons is that the two models are equally good at explaining the real brain activity. The alternative hypothesis is that the extra features in the larger model make it better at explaining brain activity; if so, then we should see statistically significant reductions in the residual gaps for the larger model, compared to the nested model.

To perform these analyses, we run spatial cluster-based permutation tests (Maris and Oostenveld, 2007) with 10,000 permutations: First, we conduct a mass-univariate one-tailed paired-samples *t*-test at all cortical sources in our ROIs. Then, we find clusters of contiguous points (sources) where the difference between models is statistically significant, and calculate the sizes of the resulting clusters (sum of absolute *t-*values within each cluster). We used threshold-free cluster enhancement (Smith and Nichols, 2009) in lieu of setting an arbitrary cutoff *α* level. To avoid arbitrarily small clusters, we only consider clusters with contributions from at least 10 sources. Next, we repeat this 10,000 times, but for each iteration, we randomly either switch or do not switch the model labels for each participant independently. The logic is that, if the null hypothesis is correct, and the two models are equally good at explaining the data, then their labels are interchangeable, and the random permutation should result in clusters that are statistically comparable in size to the real cluster. But if the alternative hypothesis is correct, then the labels matter and permuting them should result in smaller clusters. As a result, we can compare the real cluster size against a non-parametric distribution of 10,000 permuted cluster sizes. We deem the larger model a better fit than the nested model, if the real cluster is bigger than 95% (*α* =0.05) of permutation clusters. As this procedure is done independently in 4 ROIs, we also apply a False Discovery Rate (FDR) correction (Benjamini and Hochberg, 1995) to account for multiple comparisons. The output of each such spatial analysis is: statistically large clusters of neighboring cortical sources, and a *p*-value per ROI reflecting the confidence in rejecting the null hypothesis within it.

To ensure that the better fit of the larger model is truly attributable to the precise impulse timings of the extra features, and is not a random artifact of simply having extra features, we also performed a similar comparison between the larger model and three shuffled versions of itself, following (Brodbeck et al., 2018). For each shuffled model, we duplicated the real model, but randomly shifted the impulse timings of the model’s extra features within each trial. This means that the model’s extra features were not aligned with the points of interest. The null hypothesis here is that each shuffled model is equally good at explaining the cortical data as the real model, and the impulse timings of the extra features should not matter. The alternative hypothesis is that the real model is better than the shuffled models, precisely because its impulse timings are well-aligned. The procedure is the same as the regular model comparisons outlined above.

Finally, we can also visualize and statistically analyze the TRF solutions. For that, we averaged across all 20 TRF solutions per model and subject, then pooled the solutions across subjects for group analyses. For each feature of interest, and at each cortical source and time-lag (0–500 ms), we test the group-level TRF magnitude against 0 (a two-tailed one-sample *t*-test), meaning against the null hypothesis that the feature does not contribute to explaining brain activity at that cortical source and that time-lag. The procedure is similar to the one outlined above, except that here we use a *spatio-temporal* clustering algorithm, since the TRF solution is defined over space and time. Here, we also use 10,000 permutations and a cluster-size alpha level of 0.05, in addition to FDR correction to account for comparisons across ROIs. The output of this analysis for each feature is: statistically large clusters of neighboring source-time points, and a *p*-value per ROI reflecting the confidence in rejecting the null hypothesis in the ROI and time-lag window.

### Evoked response analyses

For evoked-response analyses, we aligned all MEG epochs by the point of interest (e.g., the *stem uniqueness onset*), then obtained the evoked responses per condition in sensor space, before calculating the inverse solution in the same 4 ROIs (SNR=3), morphing the results from participants’ native MRI space to the Freesurfer average surface. We then tested for group-level differences between conditions using spatio-temporal cluster-based permutation tests, similarly to the procedure outlined above. First, we performed mass-univariate two-sided paired-sample *t*-tests at each source-time point, comparing the conditions of interest; next, we applied a spatio-temporal clustering algorithm. We used cluster size (integration over local statistic values within a cluster) as a cluster-level statistic, and compared it against a null hypothesis-generated distribution of 10,000 permuted statistics. In each permutation, we chose a random subset of subjects, swapped their condition labels, and repeated the calculation above. The fraction of permuted statistics that are bigger than our real, non-permuted cluster size gave us the p-value for the tested effect within the ROI and time window of interest. Finally, we used FDR to account for multiple comparisons across the 4 ROIs.

## Results

### Behavioral results

Mean accuracy across all 27 participants and both pronoun and stem tasks was 96.89% (SD=3.28%). Mean accuracy for pronoun task items was 96.79% (SD=5.14%), and for stem task items 97.07% (SD=2.72%). There was no significant difference between task types (*p*=0.76). Since the task’s purpose was to ensure participant engagement and attention to the stimuli, we did not further analyze the behavioral data.

### BOTTOM-UP vs. NAÏVE models

In the TRF analysis, we investigated whether, compared to the NAÏVE model that is blind to morpheme information, the BOTTOM-UP model provides a better fit to the neural data. Wherever this is true, the BOTTOM-UP residual map should have smaller values than the NAÏVE model’s. The analysis revealed that the morpheme boundary feature (the only feature distinguishing the two models; Fig. 2) accounted for a significant improvement in fit to the neural data in large swaths of the bilateral temporal and inferior frontal ROIs (all *p*<0.0001; Fig. 3).

**Figure 3.**
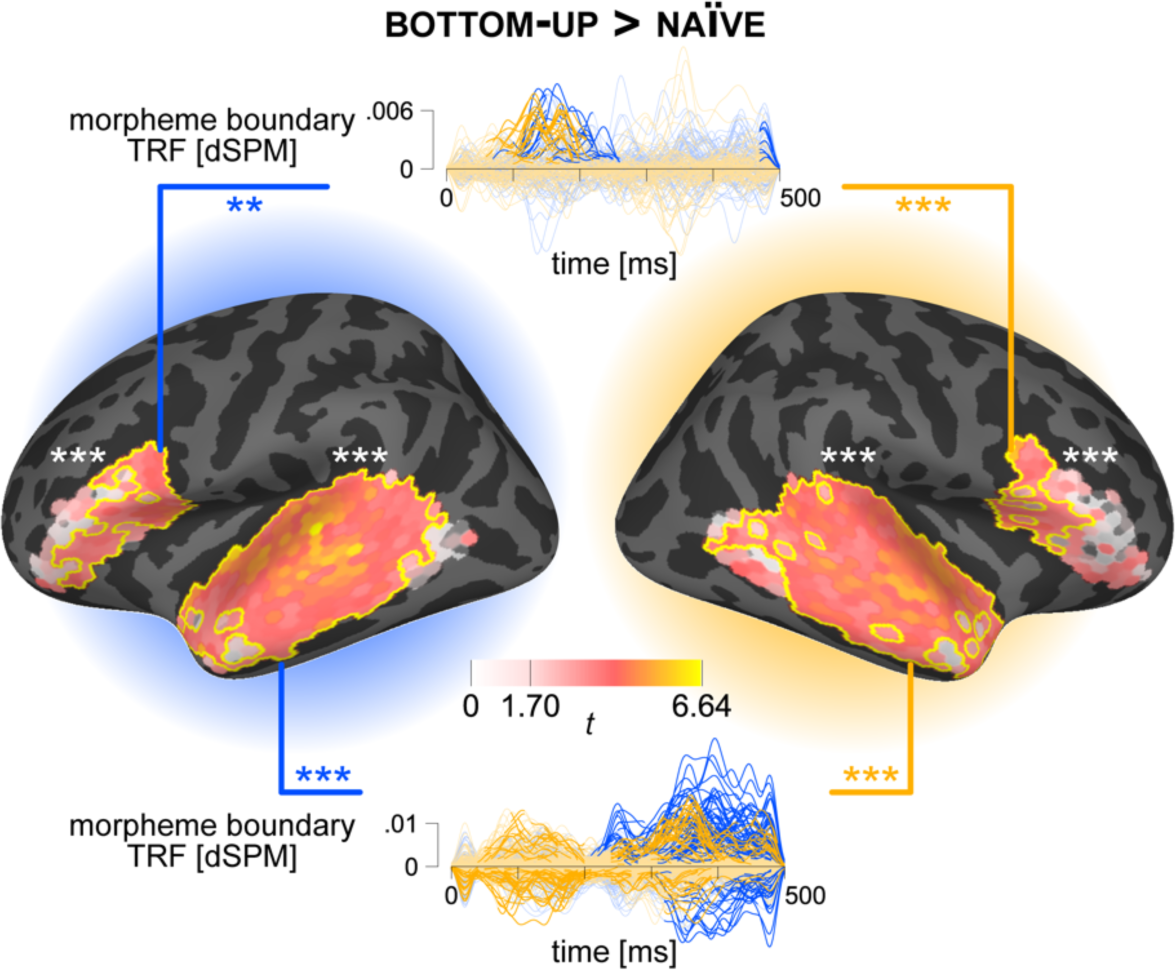
A morphologically _BOTTOM_-_UP_ model explains more fronto-temporal activity than a morphologically _NAÏVE_ model. Inflated cortical surface: masks highlight the 4 ROIs used for analysis—the bilateral temporal and inferior frontal cortices. Heat maps reflect statistical differences between the explanatory power of the two models. Each cortical source’s color reflects local t-statistic for rejecting the null hypothesis that the two models are equally good. Yellow borders indicate spatial cluster borders. White asterisks refer to *p*-values within nearby ROIs. Insets: the Temporal Response Function (TRF) solutions for the morpheme boundary feature of the _BOTTOM_-_UP_ model in the left (blue time-courses) and right (gold) frontal (top inset) and temporal (lower inset) ROIs. Each time-course corresponds to one cortical source. Highlighted sections reflect spatio-temporal clusters of a statistically significant size. Colored asterisks refer to *p*-values within corresponding ROIs and time-lag window.

To ensure that this effect is truly attributable to morpheme boundary timings, and not to the arbitrary addition of an extra impulse train to the model, we tested the BOTTOM-UP model against 3 shuffled versions of itself (see *Materials and Methods*). For each shuffled model, the impulses of the morpheme boundary feature were shifted a random amount in every trial. The real BOTTOM-UP model performed better than all 3 shuffled models across all 4 ROIs (all *p≤*0.0001; see Supplementary Fig. S2a and Supplementary Table S2).

We also tested the TRF profile of the morpheme boundary feature against 0 (see Fig. 3 insets). This revealed bilateral temporal effects (both *p*<0.0001) and bilateral inferior frontal effects (left *p*=0.0014, right *p*<0.0001). The largest clusters in the temporal ROIs spanned 70–230 ms (left) and 60–220 ms (right); in the frontal ROIs they spanned 210–500 ms (left) and 0–500 ms (right).

### PREDICTIVE models vs. BOTTOM-UP model

Next, we compared 3 PREDICTIVE morphological models against the BOTTOM-UP model. The PREDICTIVE models contained all features of the BOTTOM-UP model, plus extra morpheme-level information (see Fig. 2 and Table 1). In the bilateral temporal ROIs, all PREDICTIVE models (PATIENT, EAGER, and PROBABILISTIC) out-performed the BOTTOM-UP model (all *p*<0.0001; Fig. 4). The same was true in smaller clusters of the left inferior frontal ROI (PATIENT and PROBABILISTIC *p*<0.0001; EAGER *p*=0.0007). A right inferior frontal effect only appeared for the PATIENT PREDICTIVE model (*p*<0.0001) with a small posterior cluster (Fig. 4a). To ensure that the improvement in fit is due to the actual timings of the extra features, and not simply the addition of extra impulse trains, we compared each PREDICTIVE model with 3 shuffled versions of itself, where the extra impulses were shifted a random amount in time for each trial. The result was a better fit across the board for all real predictive models in all four ROIs, compared to their shuffled versions (see Supplementary Fig. S2b–d). Supplementary Table S2 provides the results of all the comparisons, by model, ROI, and shuffling.

**Figure 4.**
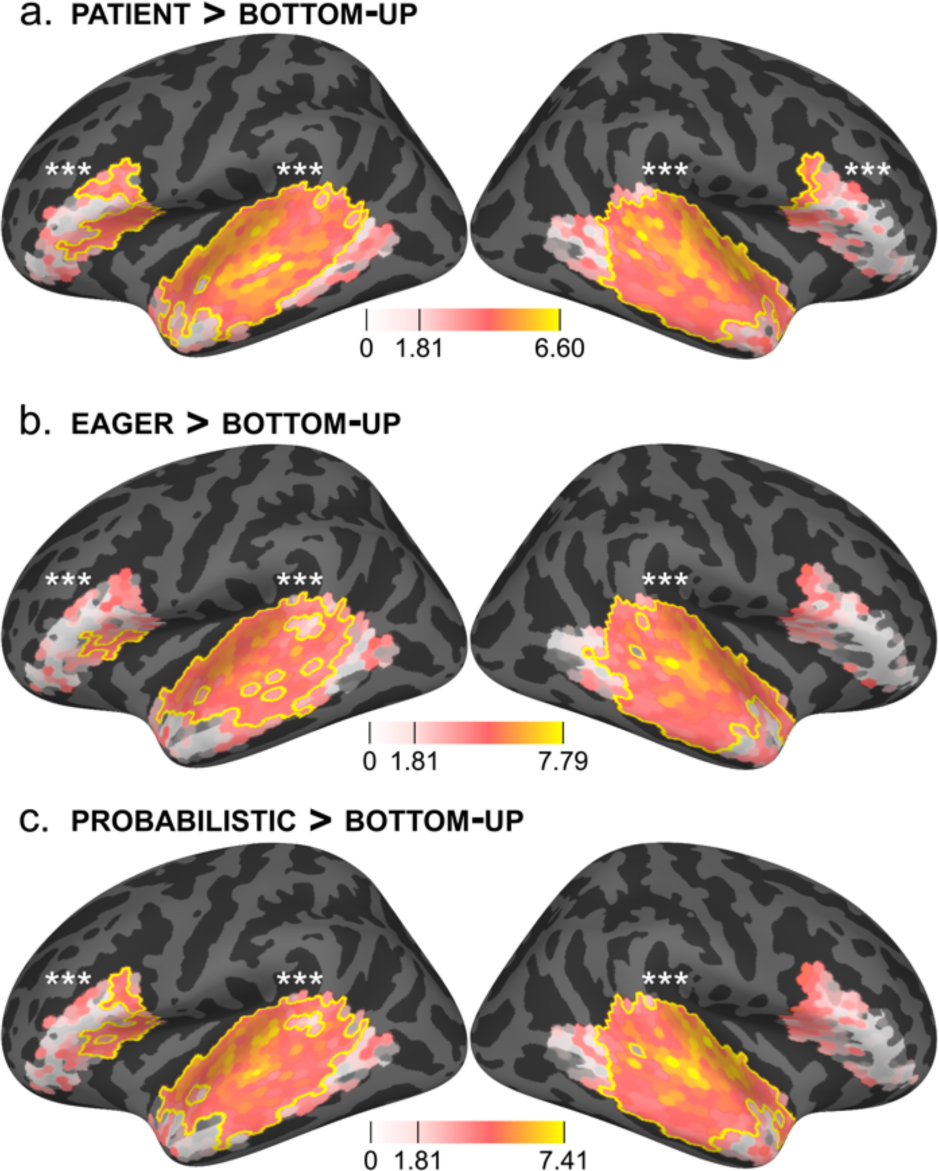
Morphologically _PREDICTIVE_ models explain more bilateral temporal and left inferior frontal activity than a morphologically _BOTTOM_-_UP_ model. Results of the statistical comparison between the explanatory power of the _BOTTOM_-_UP_ model and the (a) the _PATIENT_, (b) the _EAGER_, and (c) the _PROBABILISTIC_ models. Cortical surface: masks highlight ROIs. Heat maps reflect statistical differences between explanatory powers of model pairs. Each cortical source’s color reflects the local t-statistic for rejecting the null hypothesis. Yellow borders indicate cluster extent. White asterisks indicate *p*-values within nearby ROIs.

To better understand the contribution of the extra PREDICTIVE features, we also tested their TRF profiles. We only show temporal ROIs, as the clusters there were much larger than the frontal ROIs. The results (upsampled from 100 Hz to 1 kHz) appear in Fig. 5. Note that because 20 ms is the first boost-able time-lag (see *Materials and Methods*), we ignore peaks at 20 ms: they are implausibly early and probably indicate effects prior to the TRF time-lag window (i.e., the feature’s impulse comes ‘too late’).

**Figure 5.**
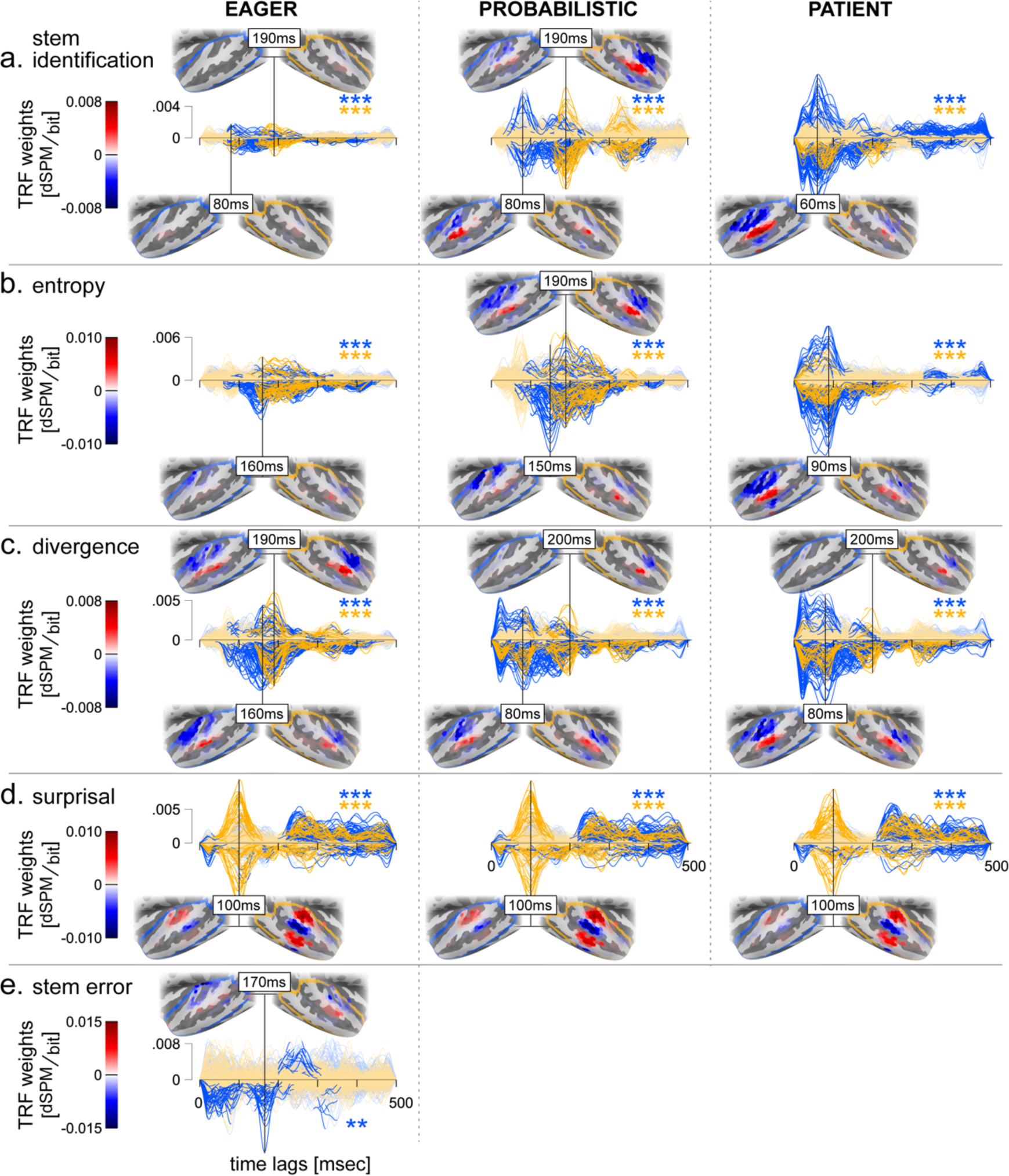
Temporal Response Functions (TRFs) of _PREDICTIVE_ morpheme-level features. Each row shows TRF solutions for the same feature: (a) stem identification, (b) morpheme entropy, (c) morphemic divergence, (d) morpheme surprisal, and (e) stem error detection (_EAGER_ only). Each column corresponds to one _PREDICTIVE_ model. Shown are TRF solutions in left (blue) and right (gold) temporal ROIs. Asterisks indicate *p*-values in left (blue) and right (gold) ROI. Each time-course corresponds to a single cortical source. Highlighted sections reflect spatio-temporal clusters. Cortical insets: cortical patterns of TRF magnitudes at indicated peak timepoints. For each feature (row), models share the same magnitude scale (for time-courses) and colorbar (for cortical patterns).

**Stem identification** (Fig. 5a) had similar TRF profiles in EAGER and PROBABILISTIC models, though the latter had a larger magnitude, peaking at a 80 ms time-lag in the left superior temporal cortex and 190 ms in its right homologue. In the PATIENT model, a left peak (and a smaller right peak) appeared earlier, at 60 ms. **Morpheme entropy** (Fig. 5b) also had similar EAGER and PROBABILISTIC profiles, again of greater magnitude in the latter, with a left superior temporal peak at a 150 ms time-lag and a right peak at 190 ms. In the PATIENT model, as before, a left peak (with a smaller right peak) appeared earlier, at 90 ms. In contrast, **morpheme divergence** (Fig. 5c) had a similar 200 ms right superior temporal TRF peak across all three models; however, an earlier left superior temporal peak appeared at 80 ms in the PROBABILISTIC and PATIENT models, though later (160 ms) in the EAGER model. The locus of **stem identification**, **morpheme entropy**, and **morpheme divergence** clusters localized to similar bilateral superior temporal regions across PREDICTIVE models.

**Morpheme surprisal** (Fig. 5d) had the same profile across models, with a large right-lateralized superior and middle temporal peak at 100 ms. The similarity across models is coherent with the fact that surprisal has impulses of almost identical timings and values across the three models (Fig. 2 and Supplementary Fig. S1). Finally, **stem error** detection (Fig. 5e) is only defined for the EAGER model, and showed a 170 ms peak corresponding to a very small superior temporal area.

In sum, all the extra features contributed to explaining some of the brain data across the three PREDICTIVE models, at differing time-lags and relative strengths in the bilateral temporal ROIs.

### PATIENT vs. PROACTIVE models

The previous results show that multiple PREDICTIVE strategies are plausible for morpheme-level speech processing. But these strategies are not mutually exclusive; the brain could hypothetically deploy and pool evidence from more than one strategy. To test this hypothesis, we performed two more nested comparisons. In both cases, the baseline is the conservative PATIENT model, in which the brain awaits complete stem disambiguation before identifying the stem. To that model, we compared two *PROACTIVE* models that process the input before disambiguation: these included the same features as the PATIENT model, plus the PREDICTIVE morpheme features of either the EAGER or the PROBABILISTIC models.

The null hypothesis here is that these EAGER+PATIENT or PROBABILISTIC+PATIENT models should be just as good as the PATIENT model at explaining the brain data, since both are PREDICTIVE and have access to similar features. The alternative hypothesis is that different strategies do not explain completely overlapping portions of the neural data, and that adding PROACTIVE (EAGER or PROBABILISTIC) features provides an even better fit to the data.

Compared to the PATIENT model, the EAGER+PATIENT model explained significantly more activity in the left and right temporal ROIs (both *p*<0.0001; Fig. 6a). The same occurs with the PROBABILISTIC+PATIENT model (both *p*<0.0001; Fig. 6b). We found no frontal effects in the two comparisons.

**Figure 6.**
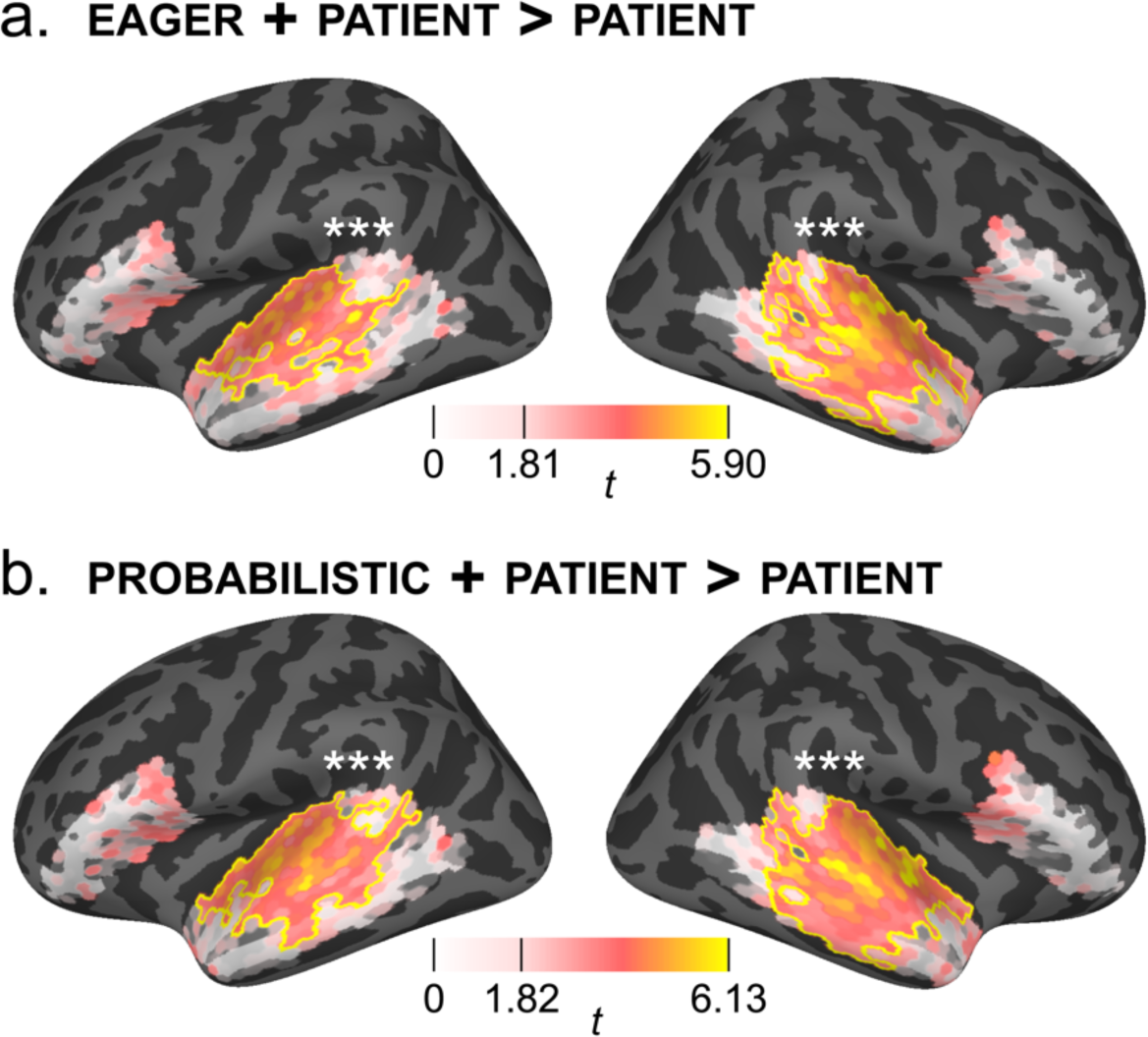
Morphologically _PROACTIVE_ strategies explain more bilateral temporal activity than a pure _PATIENT_ strategy. Results of the statistical comparison between the explanatory power of the _PATIENT_ model and 2 _PROACTIVE_ models: (a) a _PATIENT_ + _EAGER_ model, (b) and a _PATIENT_ + _PROBABILISTIC_ model. Cortical heat maps reflect the statistical difference between the explanatory power of the two models. Each cortical source’s color reflects the local t-statistic for rejecting the null hypothesis. Yellow borders indicates spatial cluster extents. White asterisks indicate *p*-values in nearby ROIs.

The TRF profiles of the extra EAGER and PROBABILISTIC features are shown in Supplementary Fig. S3. They show similar profiles to those in Fig. 5, suggesting that the differences in feature timings and values between the conservative PATIENT model and the *PROACTIVE* EAGER / PROBABILISTIC models explain different portions of the data.

### Evoked analysis

We compared cortical responses to length-UNAMBIGUOUS (‘*qayyama’*) and long length-AMBIGUOUS (‘*ṣaddaqa*’) verb stems, time-locked to *stem uniqueness onset* (see Fig. 1a). The null hypothesis is that there is no difference between how the brain processes the two conditions. Specifically, under the null hypothesis there should not be any differences between conditions either before or after *stem uniqueness* onset. This would be true of the PATIENT, BOTTOM-UP, and NAÏVE models, since none of them differentiate between the two conditions, especially as they have the same verb template (Fig. 1a).

The alternative hypothesis is that the two conditions are processed differently. This would be true of the EAGER and PROBABILISTIC models, which identify possible stems before disambiguation (Fig. 1a). In this case, we should find differences between conditions already *before* stem uniqueness onset, since a PROACTIVE brain already identifies a short stem in the length-AMBIGUOUS (‘*ṣadda*’)—but not the length-UNAMBIGUOUS—condition. We should also find differences *after* stem uniqueness onset, since a PROACTIVE brain would need to revise its stem commitment (under the deterministic EAGER strategy) or recalibrate the probabilities (under the PROBABILISTIC strategy), but only for length-AMBIGUOUS stems.

We performed the evoked analysis in the same 4 ROIs, in a time window extending from −250 ms before to 250 ms after stem uniqueness onset. We found effects in the bilateral temporal ROIs (both *p*=0.002) and the left inferior frontal ROI (*p*=0.018). *Before* stem uniqueness onset (Fig. 7a), we found clusters in the upper portion of the left (−241 to −44 ms) and right (−194 to −44 ms) superior temporal ROIs, and matching clusters of opposite polarity in the lower portion of the left (−178 to −40 ms) and right (−183 to −44 ms) superior temporal ROIs. *After* stem uniqueness onset (Fig. 7b), we found clusters in the upper portion of the left (18–132 ms) and right (49–142 ms) superior temporal ROIs, and a matching cluster in the lower portion of the superior left temporal ROI (21–132 ms). Finally, we also found a left inferior frontal cluster (−37–28 ms) straddling the stem uniqueness onset.

**Figure 7.**
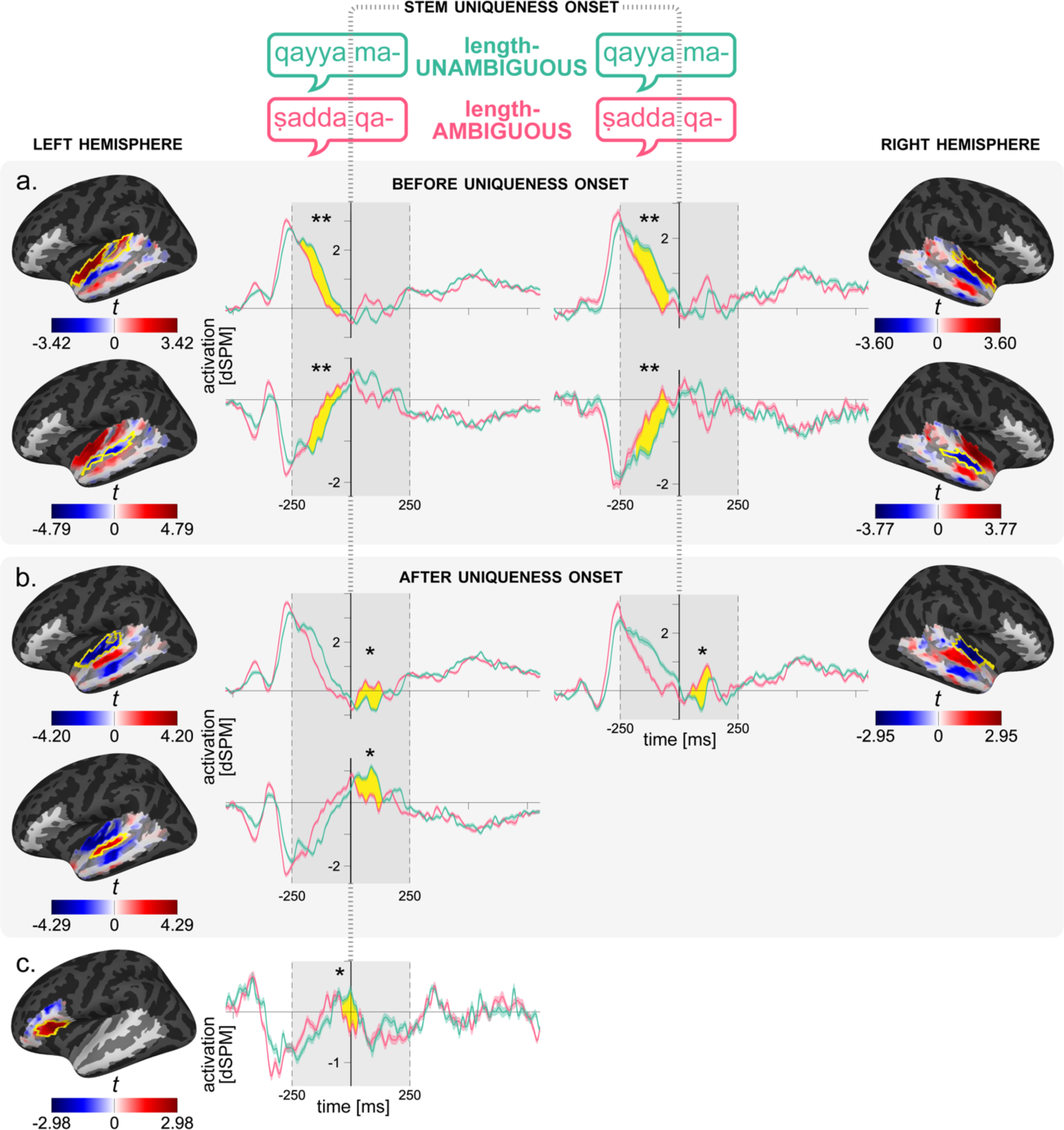
Evoked analysis reveals morphological effects before and after stem uniqueness onset. Results from the comparison of evoked source responses to long length-_AMBIGUOUS_ (pink) and _UNAMBIGUOUS_ (turquoise) items. t=0 corresponds to the stem uniqueness onset. Shown are results (a) before, (b) after, and (c) straddling stem uniqueness onset. Cortical maps: colormaps show time-averaged (within cluster) t-statistics for rejecting the null hypothesis that the two conditions are interchangeable. Yellow borders indicate spatial extents of clusters. Time-courses: evoked responses averaged over all sources that contribute at least one timepoint to corresponding clusters. Grey time window (−250–250 ms) indicates test window. Yellow fillings indicate the temporal extent of each cluster.

To ensure that these effects indeed reflect early stem identification differences between conditions, we repeated the analysis, this time aligning all trials by the onset of the second consonant (Supplementary Fig. S4). This corresponds to the *first stem identification* onset of the AMBIGUOUS condition. The analysis in a −100–250 ms window relative to this point revealed effects in the bilateral superior temporal lobe (both *p*=0.002) and the left inferior frontal ROI (*p*=0.042). The resulting clusters were very similar to the previous set of evoked results in terms of spatial patterns and temporal extents (upper left superior temporal: 59– 249 ms; lower left superior temporal: 71–239 ms; upper right superior temporal: 67–249 ms; lower right superior temporal: 77–247 ms; left inferior frontal: 201–249 ms).

## Discussion

In this speech comprehension study, we set out to investigate whether, in order to process speech, the brain tracks meaningful sub-word units called morphemes. We designed an MEG experiment using spoken Arabic sentences that were single multi-morphemic words (‘*ṣaddaqa-t-hu*’) consisting of verb stems, subject morphemes, and direct object morphemes (Fig. 1). We also proposed several hierarchically-nested encoding models that differed in whether and how they represented morpheme information (Fig. 2). Our design allowed complementary encoding model (TRF) and evoked analyses.

### Spatiotemporal dynamics of proactive and predictive spoken morpheme processing

Compared to the NAÏVE model, the BOTTOM-UP model with added morpheme boundary information explained significantly more bilateral temporal and inferior frontal activity (Fig. 3). These effects are not attributable to the mere addition of randomly-timed impulse trains (Supplementary Fig. S2a). The morpheme boundary TRFs (Fig. 3) show significant bilateral inferior frontal magnitudes in 100–200 ms time-lags, in addition to early (right) and late (bilateral) temporal contributions. The spatiotemporal patterns are in line with previous work looking at morphological effects time-locked to morpheme boundaries (Bakker et al., 2013; Whiting et al., 2013; Leminen et al., 2013; see Leminen et al., 2019, for a review).

Adding PREDICTIVE morpheme-level features (Fig. 2, Table 1) explained significantly more activity in bilateral temporal and left inferior frontal clusters (Fig. 4). This suggests that the brain predictively processes, identifies, and segments morphemes. These effects are attributable to the values and timings of the PREDICTIVE features (Supplementary Fig. S2b-d).

Only the BOTTOM-UP model explained more right frontal activity (Fig. 3; except for a small PATIENT cluster, see Fig. 4), implicating this region in bottom-up morpheme processing. In contrast, BOTTOM-UP *and* PREDICTIVE models explained more bilateral temporal and left inferior frontal activity (Figs. 3&4). The overlap in findings suggests a predictive role. For PREDICTIVE models to out-perform the BOTTOM-UP model, pre-boundary morpheme-related activity is necessary. We cannot rule out a bottom-up role here, but cannot confirm it either, since morpheme boundary and predictive TRFs overlap in time (Fig. 2). This adds to prior work implicating the region in auditory morphological processing (Tyler et al., 2005; Bozic et al., 2010).

Additionally, combining PATIENT and PROACTIVE (EAGER or PROBABILISTIC) features outperformed a PATIENT-only model in bilateral temporal ROIs (Fig. 6). This suggests that predictive strategies are not mutually exclusive, in line with evidence for the co-existence of different neural strategies generally in speech processing (Obleser and Kotz, 2011). It also implies a proactive role (e.g., either EAGER or PROBABILISTIC) in these regions, since PROACTIVE models must clear a higher (earlier) bar. But, similarly to the above logic, we cannot rule out nor confirm a PATIENT strategy there.

By contrast, adding PROACTIVE on top of PATIENT strategies did not further explain left inferior frontal activity, implicating it in PATIENT processing. However, a PATIENT account alone cannot explain the evoked effect there (Fig. 7c): it is a non-sensory area and the effect straddles uniqueness onset, suggesting it is a downstream effect of an earlier, proactive process. This implicates this region in *proactive* morpheme-level processing as well.

To sum up the spatial pattern, our findings imply proactive predictive morpheme processing in bilateral superior temporal areas, a bottom-up role for the right inferior frontal cortex, and a dual patient/proactive role in the left inferior frontal cortex.

In the time domain, results from our PATIENT model, which awaits full stem disambiguation, are congruent with behavioral work using information-theory metrics at *stem uniqueness* points (Wurm et al., 2006; Balling and Baayen, 2008, 2012). But findings from EAGER and PROBABILISTIC models, which are PROACTIVE*—*processing morpheme information before stem disambiguation*—* show the relevance of an earlier critical point: the *first stem identification* point (Fig. 1a), at which the accumulating input first forms an existing stem.

Our evoked analyses strengthen this point: they revealed differences between responses to length-AMBIGUOUS and length-UNAMBIGUOUS stems in the bilateral superior temporal cortex, both *before* and *after* stem uniqueness onset (Fig. 7a&b). A follow-up analysis showed that these clusters appear ∼70 ms after early identification onset (Supplementary Fig. S4). These results provide further evidence for a neural relevance to the *first stem identification* point, and for a *proactive* account, since only a brain that can use a PROACTIVE strategy can tell the two conditions apart *before* and *after* stem uniqueness onset.

### Spatiotemporal dynamics of predictive morpheme-level features

We also tested the spatiotemporal TRF profiles of PREDICTIVE features (Fig. 5). To our knowledge, very little work has published neural results from morphemic information-theory metrics, time-locked to morphologically crucial time-points. Thus, where unavailable, we compare effect timings to other levels of predictive speech processing.

Morpheme surprisal elicited an early ∼100 ms peak in the superior and middle temporal cortex (Fig. 5d) across strategies. (Surprisal is almost identically defined across models; Fig. 2, Table 1). (Gwilliams and Marantz, 2015) found morpheme surprisal effects in the left superior temporal cortex between 200–400 ms, similar to our later left-hemisphere results; however, they only analyzed left-hemispheric activity. Our ∼100 ms peak timing is similar to phoneme surprisal timings from past studies (Brodbeck et al., 2018, 2023; Donhauser and Baillet, 2020). Though phoneme surprisal peaks are bilateral, our main peak is right-biased, suggesting a distinction between phoneme- and morpheme-surprisal.

The remaining predictive features’ TRFs differed across models, but generally localized to the bilateral superior temporal cortex. For stem identification (Fig. 5a) and morpheme entropy (Fig. 5b), PROBABILISTIC TRFs were enhanced versions of EAGER TRFs, with similar peaks. In contrast, PATIENT TRFs were distinct and peaked earlier. Entropy peaks are also similar in timing to previous phoneme entropy findings (Brodbeck et al., 2018, 2023; Donhauser and Baillet, 2020). As to divergence (Fig. 5c), PROBABILISTIC and PATIENT TRFs were similar, with EAGER TRFs distinct. Finally, the EAGER model had a unique stem error feature, whose TRFs peak at 170 ms in a small superior temporal cluster.

The TRF results suggest that during predictive processing, these features engage the brain at different time-points. They also show that the PROBABILISTIC solution is a hybrid of the risk-averse PATIENT solution and the risk-taking EAGER solution; this is unsurprising, since the PROBABILISTIC strategy is a weighted average of the other two. Across predictive features, the PROBABILISTIC solutions are the most diverse, suggesting that its different features explain distinct portions of the brain data.

In contrast, PATIENT peaks, which are time-locked to *stem uniqueness onset*, may be implausibly early; indeed, most peaked at 20 ms—the very first boost-able time-lag that maintains a causal solution (see *Materials and Methods*). This hints at feature-related activity already before *stem uniqueness onset* (t=0 in PATIENT model): the PATIENT model alone may be too late in explaining neural activity. This reinforces our conclusions from Fig. 6 & Fig. 7.

### Implications

Our results lend neural support to the role of the stem uniqueness point (Balling and Baayen, 2008) during speech processing. However, they also offer an earlier, equally important point: the *first stem identification* point—the first point at which any stem is identifiable, even if not uniquely. Note that these are likely not the only morphemically relevant point. For example, Arabic stems are themselves decomposable into roots and templates (Fig. 1a), and (Gwilliams and Marantz, 2015) have shown root-related effects in auditory word comprehension. Here, we isolated stem-level computations from stem-internal ones by using the same verb templates across items.

The real neural algorithms are likely to be more complex than our models, especially when considering natural language. For example, consider the two phrases ‘*rain jacket*’ and ‘*range accuracy*’. In connected speech, both have the same onset /reɪnʤækə/ in American English. A PATIENT strategy as proposed here would initially identify the stem ‘*range*’ for both cases, necessitating an error-correction mechanism to parse ‘*rain jacket*’. A PROACTIVE strategy as proposed here initially identifies ‘*rain*’ for both phrases. Then, it either hyper-corrects both to ‘*range*’, or sticks to ‘*rain*’ but fails to find any stem congruent with /dʒækərə/ in ‘*range accuracy*’, forcing it to step back. Future work on naturalistic speech processing may need to refine these models to combine strategies, include more decision points, and possibly add a stack of recently identified but revisable morphemes, in addition to top-down contextual constraints.

Our findings highlight a critical gap in many neural and computational models of speech comprehension, which typically include word- and phoneme-level information, but exclude morpheme information. We show that accounting for morphemes significantly improves models’ abilities to explain neural data.

Our findings also add to previous work showing temporal cortex involvement in predictive morphological processing during speech comprehension (Ettinger et al., 2014; Gwilliams and Marantz, 2015). More generally, our findings lend support to predictive models of speech and language processing, extending them to the morpheme level.

Finally, to our knowledge, this is the first study investigating morpheme-level processing in full, grammatical sentences. Rather than spoken word frameworks or violation paradigms, then, our results are interpretable within and contribute to the sentence processing literature. Particularly, they mirror similar tensions between probabilistic and deterministic syntactic parsing strategies (Crocker, 1999).

## Conclusion

In this speech comprehension study, we investigated whether and how the brain processes sub-word meaning-carrying linguistic units, called morphemes. To that end, we used MEG to measure brain responses to short multi-morphemic sentences and compared the fit of hierarchically nested cognitive models to the brain data. We showed that, above and beyond acoustic, phonetic, syllabic, and lexical information, adding morpheme-level information to the models explains more data. Specifically, adding morpheme boundary information explains more data in the bilateral temporal and inferior frontal cortices, while adding predictive morphemic features explains more data mainly in the bilateral superior temporal cortices, and in the left inferior frontal cortex. Using two complementary analyses, we also show evidence of the brain using PROACTIVE predictive strategies, which process accumulating spoken morphemes even before they are fully disambiguated. Our results highlight the importance of updating existing speech comprehension models to include morpheme-level processes.

## Acknowledgments

This research was supported by the NYUAD Research Institute under Grant G1001, the Economic and Social Research Council (ESRC) in the United Kingdom [ES/V000012/1], and the IKUR NEUROBIOSCIENCES 2021–2022 Research Project. This work was supported in part through the NYU IT High Performance Computing resources, services, and staff expertise.

