## Supplementary figures and tables for "Neural bases of proactive and predictive processing of meaningful sub-word units in speech comprehension"

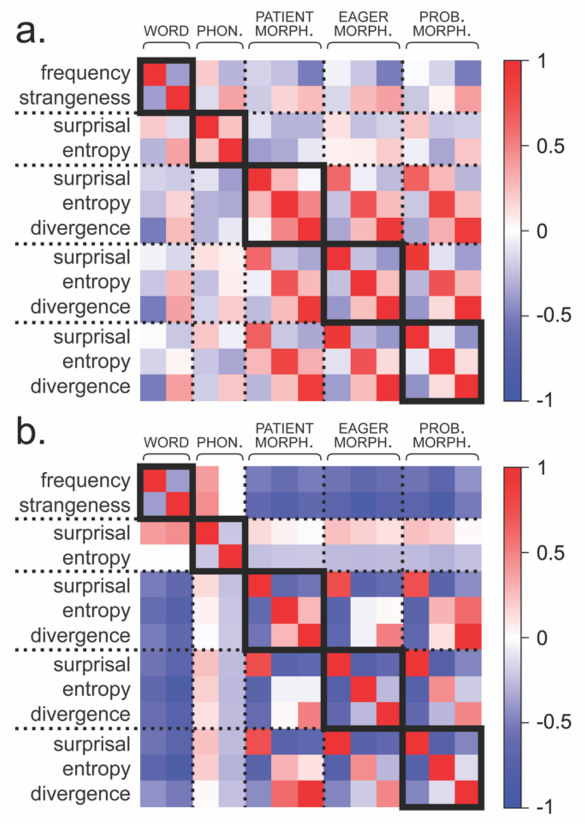


**Fig. S1. Feature correlations.** Shown are Pearson correlations (colorbar) between impulse features within and across levels of linguistic hierarchy, calculated in two ways. (a) average impulse value correlations: for each feature, its impulse values are averaged per trial, and all pairs of resulting vectors are correlated. (b) impulse timing and value correlations: for each pair of features, we extract their values at all time-points in which at least one feature in the pair is non-zero and correlate those values. Dashed lines separate between levels of linguistic hierarchy or predictive strategy. Bold squares indicate features within the same family.

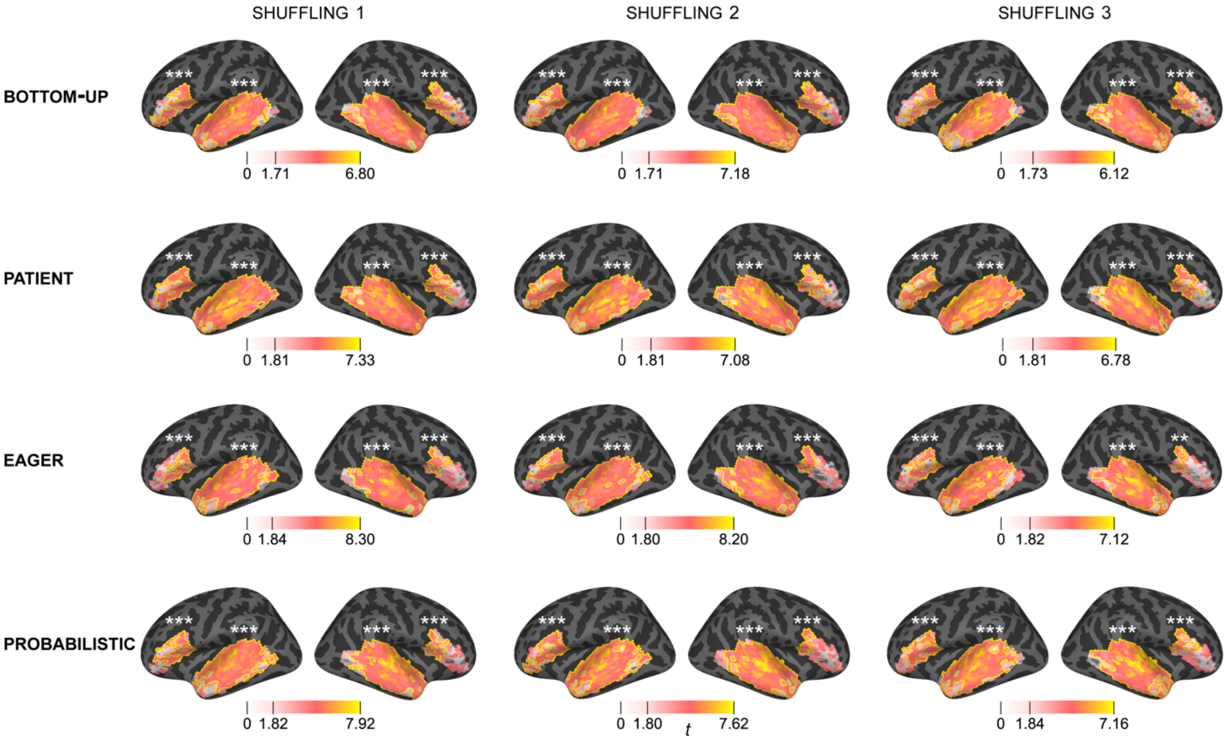


**Fig. S2. Real vs. shuffled model comparisons**. Each row shows the results of a comparison between the indicated model and three shuffled versions of itself (see *Materials & Methods*). The shuffled versions are duplicates of the original models, except that the features of interest (BOTTOM-UP: morpheme boundary; PATIENT, EAGER, and PROBABILISTIC: all predictive features) have impulses that are randomly shuffled in time per trial. Cortical heat maps reflect the statistical difference between the explanatory power of the two models. Each cortical source's color reflects the local $t$-statistic for rejecting the null hypothesis that the two models compared are equally good at explaining the source's activity. The yellow border indicates the extent of the spatial cluster identified. White asterisks indicate $p$-values within nearby ROIs.

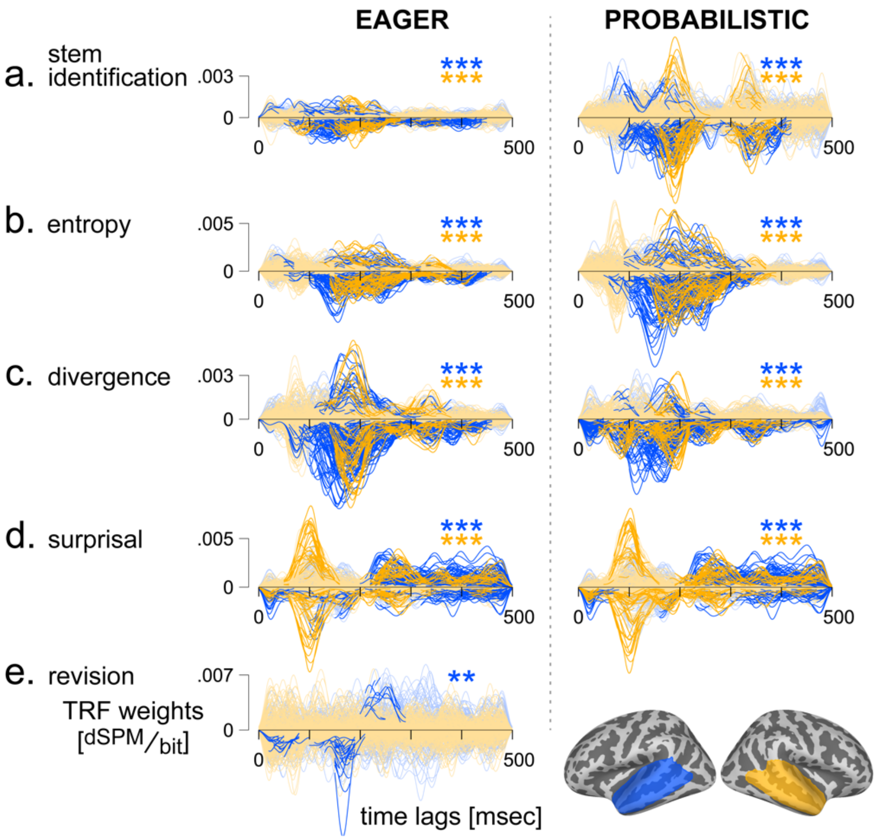


**Fig. S3. Temporal Response Functions (TRFs) of PREDICTIVE features in PROACTIVE models**. Shown are the TRF solutions in the left (blue) and right (gold) temporal ROIs. Asterisks indicate *p*-values in left (blue) and right (gold) ROIs. Each time-course corresponds to a single cortical source. Highlighted sections indicate spatio-temporal clusters. For each feature (row), models share the same magnitude scale. The stem error dete€on (e) is only defined for the EAGER + PATIENT model.

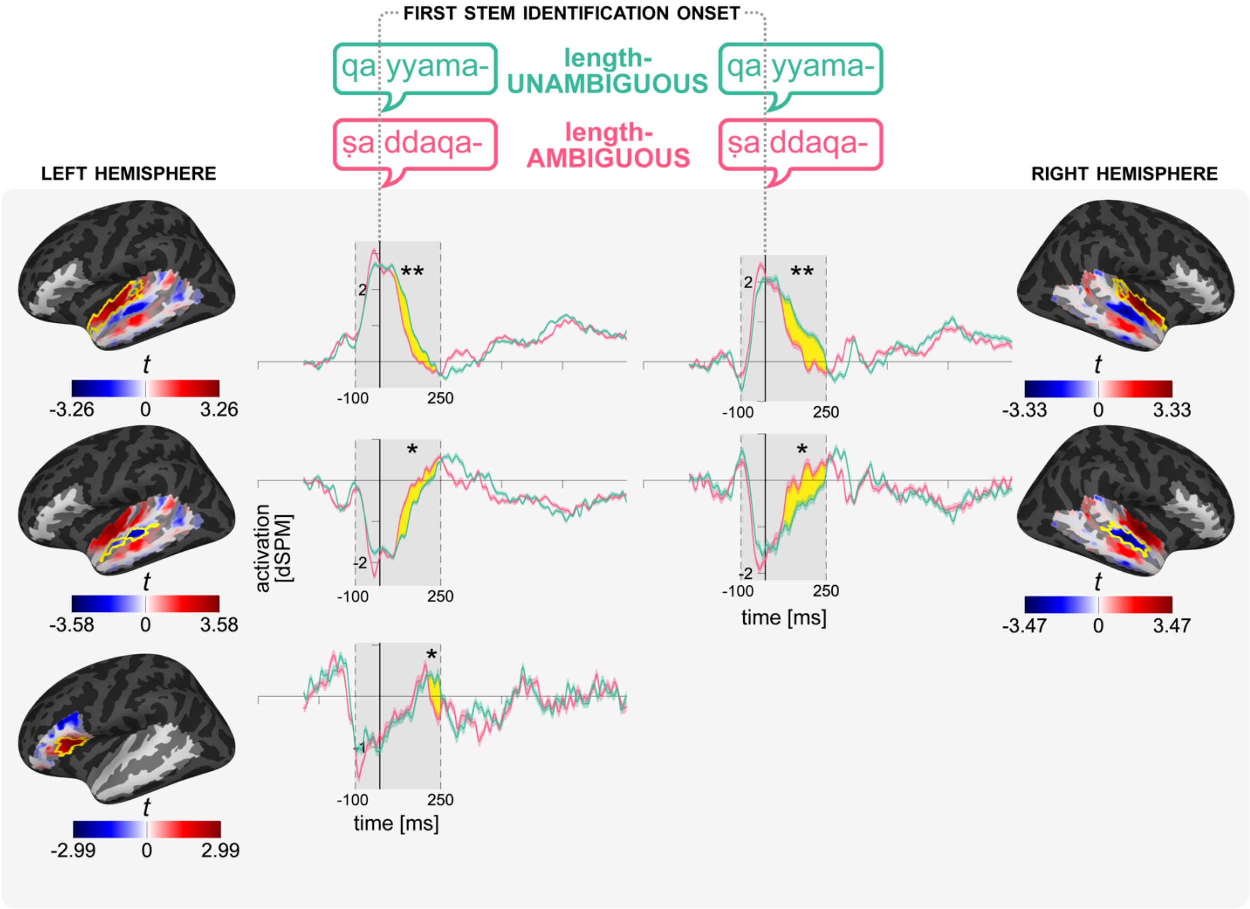


**Fig. S4 Evoked analysis to the early *first stem identification* onset.** Results from the comparison of evoked source responses to long length-AMBIGUOUS (pink) and UNAMBIGUOUS (turquoise) items. t=0 corresponds to the onset of the second consonant, which is the *first identification onset* in the AMBIGUOUS condition..Cortical maps: colormaps show time-averaged (within cluster) t-statistics for rejecting the null hypothesis that the two conditions are interchangeable. Yellow borders indicate spatial extents of clusters. Time-courses: evoked responses averaged over all sources that contribute at least one timepoint to corresponding clusters. Grey time window (-100–250 ms) indicates test window. Yellow fillings indicate the temporal extent of each cluster.

**Supplementary Table 1. Stems used as experimental material**. 30 items each for short length-AMBIGUOUS stems, long length AMBIGUOUS stems, and length-UNAMBIGUOUS stems.

| Length-AMBIGUOUS stems (short) | | | Length-AMBIGUOUS stems (long) | | |
|---|---|---|---|---|---|
| Translation | Arabic | Romanization | Romanization | Arabic | Translation |
| guided | دَلَّ | dalla | dallala | دَلَّلَ | pampered |
| cut | قَصَّ | qaṣṣa | qaṣṣara | قَصَّرَ | shortened |
| lost | ضَلَّ | ḍalla | ḍallala | ضَلَّلَ | misled |
| sprayed | رَشَّ | rašša | raššaḥa | رَشَّحَ | nominated |
| loved | حَبَّ | ḥabba | ḥabbaba | حَبَّبَ | endeared |
| pumped | ضَخَّ | ḍaḫḫa | ḍaḫḫama | ضَخَّمَ | enlarged |
| extended | مَدَّ | madda | maddada | مَدَّدَ | stretched |
| blocked | صَدَّ | ṣadda | ṣaddaqa | صَدَّقَ | believed |
| unscrewed | فَكَّ | fakka | fakkaka | فَكَّكَ | disassembled |
| dragged | جَرَّ | jarra | jarraba | جَرَّبَ | tried |
| singled out | خَصَّ | ḫaṣṣa | ḫaṣṣaṣa | خَصَّصَ | dedicated |
| settled | فَضَّ | faḍḍa | faḍḍala | فَضَّلَ | preferred |
| demolished | هَدَّ | hadda | haddada | هَدَّدَ | threatened |
| felt | جَسَّ | jassa | jassada | جَسَّدَ | embodied |
| pulled | شَدَّ | šadda | šaddada | شَدَّدَ | emphasized |
| doubted | شَكَّ | šakka | šakkala | شَكَّلَ | formed |
| cleaved | شَقَّ | šaqqa | šaqqaqa | شَقَّقَ | cracked |
| crammed | رَصَّ | raṣṣa | raṣṣaʿa | رَصَّعَ | inlaid |
| solved | حَلَّ | ḥalla | ḥallala | حَلَّلَ | analyzed |
| broke into pieces | فَتَّ | fatta | fattaša | فَتَّشَ | searched |
| cursed | سَبَّ | sabba | sabbaba | سَبَّبَ | caused |
| filled | عَمَّ | ʿamma | ʿammama | عَمَّمَ | spread |
| delighted | سَرَّ | sarra | sarraba | سَرَّبَ | leaked |
| counted | عَدَّ | ʿadda | ʿaddala | عَدَّلَ | straightened |
| replied | رَدَّ | radda | raddada | رَدَّدَ | repeated |
| landed | حَطَّ | ḥaṭṭa | ḥaṭṭama | حَطَّمَ | destroyed |
| shut | سَدَّ | sadda | saddada | سَدَّدَ | aimed |
| liked | وَدَّ | wadda | waddaʿa | وَدَّعَ | bid farewell |
| delimited | حَدَّ | ḥadda | ḥaddada | حَدَّدَ | determined |
| dipped | غَطَّ | ġaṭṭa | ġaṭṭasa | غَطَّسَ | submerged |
| | | | Length-unambiguous (long) | | |
| | | | Romanization | Arabic | Translation |
| | | | fawwaḍa | فَوَّضَ | delegated |
| | | | ġayyara | غَيَّرَ | changed |
| | | | ṣawwara | صَوَّرَ | photographed |
| | | | ḫayyara | خَيَّرَ | made choose |
| | | | ṭawwaqa | طَوَّقَ | surrounded |
| | | | qayyama | قَيَّمَ | evaluated |
| | | | zawwada | زَوَّدَ | supplied |

| | | | | | |
|---|---|---|---|---|---|
| | | | qayyada | قَيَّدَ | tied |
| | | | ḥawwala | حَوَّلَ | transformed |
| | | | ḍayyaʿa | ضَيَّعَ | lost |
| | | | ʿawwaḍa | عَوَّضَ | made up for |
| | | | mayyaza | مَيَّزَ | distinguished |
| | | | ḫawwafa | خَوَّفَ | frightened |
| | | | hayyaʾa | هَيَّأَ | prepared |
| | | | mawwala | مَوَّلَ | financed |
| | | | ġayyaba | غَيَّبَ | concealed |
| | | | ʿawwada | عَوَّدَ | habituated |
| | | | ʿayyaša | عَيَّشَ | sustained |
| | | | ḏawwaba | ذَوَّبَ | melted |
| | | | šayyada | شَيَّدَ | built |
| | | | ṭawwara | طَوَّرَ | developed |
| | | | sayyara | سَيَّرَ | pushed along |
| | | | lawwaṯa | لَوَّثَ | polluted |
| | | | dawwaḫa | دَوَّخَ | made dizzy |
| | | | šawwaqa | شَوَّقَ | fascinated |
| | | | dawwara | دَوَّرَ | recycled |
| | | | rawwaḍa | رَوَّضَ | tamed |
| | | | ʿayyara | عَيَّرَ | rebuked |
| | | | ṭawwaʿa | طَوَّعَ | enlisted |
| | | | ṣawwaba | صَوَّبَ | aimed |

**Supplementary Table S2**. ***p*-values for comparisons between models and shuffled models**. Values reflect the probability of accepting the null hypothesis that the real and shuffled models are equally good at explaining neural data. See Supplementary Fig. S2 for spatial cluster results.

| Model | ROI | Shuffling 1 | Shuffling 2 | Shuffling 3 |
|---|---|---|---|---|
| BOTTOM-UP | Left frontal | < 0.0001 | < 0.0001 | 0.0001 |
| | Right frontal | < 0.0001 | < 0.0001 | < 0.0001 |
| | Left temporal | < 0.0001 | < 0.0001 | < 0.0001 |
| | Right temporal | < 0.0001 | < 0.0001 | < 0.0001 |
| PATIENT | Left frontal | < 0.0001 | < 0.0001 | < 0.0001 |
| | Right frontal | < 0.0001 | < 0.0001 | < 0.0001 |
| | Left temporal | < 0.0001 | < 0.0001 | < 0.0001 |
| | Right temporal | < 0.0001 | < 0.0001 | < 0.0001 |
| EAGER | Left frontal | 0.0005 | 0.0007 | 0.0001 |
| | Right frontal | 0.0003 | 0.0001 | 0.0052 |
| | Left temporal | 0.0002 | 0.0001 | 0.0001 |
| | Right temporal | 0.0002 | 0.0001 | 0.0001 |
| PROBABILISTIC | Left frontal | 0.0001 | < 0.0001 | 0.0001 |
| | Right frontal | 0.0001 | 0.0001 | < 0.0001 |
| | Left temporal | < 0.0001 | < 0.0001 | < 0.0001 |
| | Right temporal | < 0.0001 | < 0.0001 | < 0.0001 |
